# Mapping Large-scale Spatiotemporal Dynamics of Synaptic Plasticity and LTP for Memory Encoding in the Hippocampal Network

**DOI:** 10.1101/2024.05.23.595474

**Authors:** Shahrukh Khanzada, Xin Hu, Brett Addison Emery, Władysław Średniawa, Daniel K Wójcik, Gerd Kempermann, Hayder Amin

**Affiliations:** Group of “Biohybrid Neuroelectronics”, German Center for Neurodegenerative Diseases (DZNE), Tatzberg 41, 01307, Dresden, Germany; Group “Adult Neurogenesis”, German Center for Neurodegenerative Diseases (DZNE), Dresden, Germany; Center for Regenerative Therapies TU Dresden (CRTD), Fetscherstraße 105, 01307, Dresden, Germany; Laboratory of Neurophysiology of Mind, Centre of Excellence for Neural Plasticity and Brain Disorders (BrainCity), Nencki Institute of Experimental Biology of Polish Academy of Sciences, 3 Pasteur Street, Warsaw 02-093, Poland; Laboratory of Neuroinformatics, Nencki Institute of Experimental Biology of Polish Academy of Sciences, 3 Pasteur Street, Warsaw 02-093, Poland; TU Dresden, Faculty of Medicine Carl Gustav Carus, Bergstraße 53, 01069, Dresden, Germany

**Keywords:** Network-LTP, Synaptic plasticity and memory, HD-CMOS-MEA, Computational dynamics, Cell assemblies, Hippocampal Network

## Abstract

Understanding memory formation requires elucidating the intricate dynamics of neuronal networks in the hippocampus, where information is encoded and processed through specific activity patterns and synaptic plasticity. Here, we introduce “EVOX,” an advanced network electrophysiology platform equipped with high-density microelectrode arrays to capture critical network-level synaptic dynamics integral to learning and memory. This platform surpasses traditional methods by enabling label-free, high-order mapping of neural interactions, providing unprecedented insights into network Long-Term Potentiation (LTP) and evoked synaptic transmission within the hippocampal network. Utilizing EVOX, we demonstrate that high-frequency stimulation induces network-wide LTP, revealing enhanced synaptic efficacy in previously inactive cell assemblies in hippocampal layers. Our platform enables the real-time observation of network synaptic transmission, capturing the intricate patterns of connectivity and plasticity that underpin memory encoding. Advanced computational techniques further elucidate the mesoscale transmembrane generators and the dynamic processes that govern network-level memory encoding mechanisms. These findings uncover the complex dynamics that underlie learning and memory, showcasing EVOX’s potential to explore synaptic and cellular phenomena in aging circuits. EVOX not only advances our understanding of hippocampal memory mechanisms but also serves as a powerful tool to investigate the broader scope of neural plasticity and network interactions in healthy and diseased states.

## 1 Introduction

Memory, a pivotal cognitive function, is more than just a repository of our acquired information. It is a dynamic neurobiological construct that intertwines with our perceptions, behaviors, and future anticipations^1^. Learning and memory involve a cascade of molecular, cellular, and neural circuit events that encode, store, consolidate, and retrieve experiences, shaping our cognitive and behavioral repertoire^2^. Like a kaleidoscope creating patterns through mirror rotations and glass arrangements, memories are believed to be represented by coordinated activity patterns through the interaction of cell assemblies across distributed brain networks during specific experiences and behaviors^3–5^. Despite extensive research, the emergence of memories through large-scale neuronal interactions remains elusive. Understanding the computational dynamics of memory mechanisms can illuminate cognitive capacities and facilitate interventions for memory disorders. While multiple brain regions contribute to various aspects of learning and memory, the hippocampus is central to spatial and episodic contexts, with its unique architecture and specialized cell types enabling the encoding of experiences into long-lasting memory traces^6–8^. Within the hippocampal circuit, information flows through interconnected regions - the dentate gyrus (DG), CA3, CA2, and CA1 —each contributing uniquely to the processing and storage of memories^6,9^. The transmission is facilitated by pathways like the perforant path, mossy fiber, and Schaffer collateral, which are modulated by synaptic plasticity^10^. Activity-dependent synaptic plasticity, particularly long-term potentiation (LTP), has been identified as a cellular substrate for learning and memory, involving intricate presynaptic and postsynaptic modifications^11^.

LTP is categorized by persistent enhancement in synaptic transmission following high-frequency stimulation, while long-term depression (LTD) is characterized by a durable decrease in synaptic strength. LTP and LTD modulate synaptic connections within ensembles, enabling efficient signal transmission essential for memory encoding, consolidation, and recall^12^. This modulation, orchestrated by synaptic plasticity, allows for the formation and stabilization of specific sequential firing patterns of particular cell assemblies, embodying the physical substrate of memory engrams^13,14^.

LTP and synaptic plasticity have been widely investigated using extracellular field recordings and conventional microelectrode array (MEA) approaches^15,16^. This has been significantly enriched by the employment of brain slice models, providing a controllable environment to dissect the cellular and network mechanisms underlying these phenomena^17^. Despite the invaluable insights provided by these methodologies, several limitations have emerged. This includes the lack of spatial resolution to accurately map synaptic activity-dependent changes and significant inter-experimental variability that hinder the reproducibility of findings and mask subtle alterations in synaptic plasticity and LTP under different experimental conditions. These limitations have underscored the need for novel large-scale recording techniques that will not only enhance our understanding of the cellular and network mechanisms underlying LTP but also provide a comprehensive spatiotemporal dynamic view of cell assemblies computations in different hippocampal subfields underlying learning and memory processes^18–20^.

Advanced electrophysiological recording techniques using high-density CMOS-MEAs have revolutionized our understanding of multimodal neural dynamics. These platforms allow simultaneous recording from multiple sites at high spatiotemporal resolution, providing a holistic view of neural interactions^20–24^. In this study, we present EVOX, a high-density, high-resolution electrophysiological platform that enables multi-site, long-term, and label-free simultaneous measurements of network-LTP and extracellular evoked synaptic potentials in the hippocampal circuit. By integrating large-scale neural recordings (i.e., 4096-microelectrodes), advanced computational tools, and a deep understanding of neural dynamics, this study aims to map the reconfiguration of large-scale cell assemblies underlying learning processes and memory encoding through studying network-wide LTP and synaptic plasticity.

Our platform facilitates stimulation and recording of electrically-induced synaptic responses from two hippocampal canonical pathways - medial perforant path (mPP) to the DG and Schaffer collaterals (SC) from CA3 to CA1. It aligns cellular layers of network-induced synaptic responses with their anatomical counterparts within hippocampal subfields using automated spatial and temporal waveform-based classification. These field-excitatory-post-synaptic potentials (fEPSPs) and the repertoire of induced synaptic responses are identified simultaneously from a group of cell assemblies encoded in firing electrodes in DG, CA3, and CA1. Our network-wide recordings also enable detailed computation of neuronal transmembrane current sources and sink generators using the kernel current source density (kCSD) method^25^. Utilizing EVOX, we evaluated network-wide evoked representations in aging hippocampal circuits, revealing how deactivated cell assemblies contribute to functional remodeling ^26^. Our findings indicate that aging impacts synaptic plasticity by reducing the activity of specific firing cells in specific layers, leading to alterations in large-scale memory-coding networks. This study provides insights into how age-related changes in synaptic efficacy and connectivity affect the overall organization and functionality of hippocampal circuits.

This is the first report on label-free, large-scale mapping of functional synaptic activity-dependent changes and network-LTP induction modulated by electrical stimulation in the hippocampus. While primarily aimed at understanding learning and memory with unprecedented spatiotemporal resolution, our research has broader implications. The platform paves the way for advancements in neuroscience, technology, artificial intelligence, and adaptive learning systems, potentially revolutionizing lifelong learning machines, neuromorphic computing, and brain-inspired machine-learning algorithms ^27,28^.

### 2 Results

#### 2.1 Implementation of the EVOX platform

To comprehensively investigate synaptic transmission and LTP across the intricately hippocampal layers, we developed the Evoked Network-wide Synaptic Potential Explorer (EVOX). This platform combines advanced experimental-computational techniques (**Figure 1**). Central to EVOX is a high-density planar CMOS-based microelectrode array (HD-CMOS-MEA) with 4096 microelectrodes^22,23^, offering unprecedented spatial and temporal resolution in recording field-evoked potential responses from the entire hippocampal circuitry. For precisely delivering electrical stimuli to the neural tissue, we use external platinum bipolar stimulating electrodes connected through a specialized input-output connector to minimize stimulation artifacts. A zero-drift triple-axis micromanipulator ensures precise and stable electrode positioning, while a custom-designed stereomicroscope aligns hippocampal tissue with the CMOS-MEA, optimizing recorded signal fidelity and accuracy (**Figure 1a**). The platform measures simultaneous network-wide evoked activity in the CA1-CA3 and DG circuits by stimulating the Schaffer Collateral (SC) and medial perforant pathways (mPP). This dual pathway stimulation can be conducted using two external electrodes or targeted stimulation within a specific CA1-CA3 or DG network (**Figure 1b**). Further enhancing the platform’s capabilities is an integrated Python-based computational framework to process and analyze the extensive multidimensional datasets, extracting spatiotemporal features such as network potentiation maps, multi-layer waveforms, and clustering algorithms. It includes tools for frequency-time dynamics, kernel current-source density (kCSD) analysis, and various statistical metrics, providing a comprehensive understanding of real-time network dynamics rooted in synaptic plasticity (**Figure 1c**). The advanced capabilities of the EVOX platform enable a nuanced exploration of synaptic dynamics, setting the stage for a detailed investigation into how these dynamics underpin network-wide LTP and its role in memory encoding, as we will explore in the following section.

**Figure 1.**
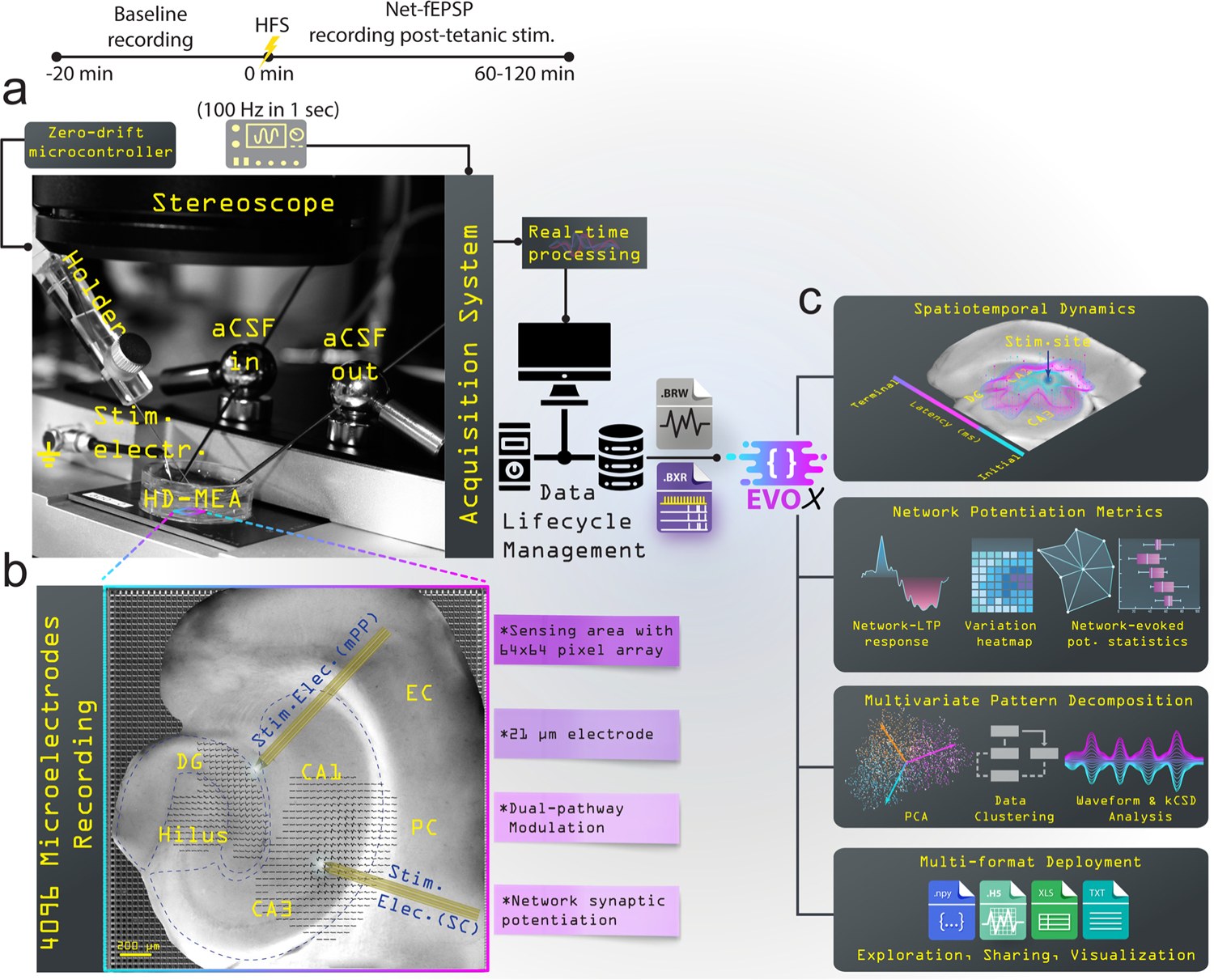
Comprehensive Overview of the EVOX Platform for Network-Level Synaptic and LTP Analysis. **a)** The platform facilitates advanced synaptic transmission and network-LTP investigation across hippocampal layers using HD-CMOS-MEA with 4096 microelectrodes. The setup features a zero-drift triple-axis micromanipulator system for precise stimulating electrode positioning and a custom-designed stereomicroscope for accurate alignment with hippocampal tissue. The experimental workflow includes a timeline from baseline measurement to tetanic-evoked potentiation achieved with a single high-frequency stimulation (HFS) at 100 Hz. **b)** The system can stimulate the Schaffer Collateral (SC) and medial perforant pathways (mPP) to measure simultaneous network-wide evoked activity, capturing distinct EPSP and PS signatures across CA1-CA3 and DG layers using bipolar stimulating electrodes. **c)** The integrated Python-based computational framework processes and visualizes the extensive multidimensional data, extracting unique spatiotemporal features, generating network potentiation metrics and multi-layer waveforms, and performing clustering and multivariate pattern analysis. This platform offers flexible, multi-format deployment options for in-depth exploration, data sharing, and visualization.

#### 2.2 Characterizing Sequential Encoding Patterns of Network-wide LTP

A key feature of EVOX is integrating functional bioelectrical network data onto corresponding optical images of the hippocampal subregions. This allows for precise mapping of evoked synaptic activation in various neuronal populations, including pyramidal cells and interneurons across distinct hippocampal layers, which is crucial for understanding the mechanisms of synaptic plasticity in memory formation^29^. This detailed mapping spanned multiple layers, including the stratum oriens (SO), stratum pyramidale (SP), stratum radiatum (SR), and stratum lacunosum-moleculare (SLM) in the CA1-CA3 network, as well as the molecular layer (ML), granule cell layer (GCL) and the Hilus (H) in the DG network^30^ (**Figures 2a,b**). Understanding how sequential encoding patterns of network-wide LTP contribute to hippocampal learning and memory is crucial for elucidating the underlying mechanisms of synaptic plasticity.

**Figure 2.**
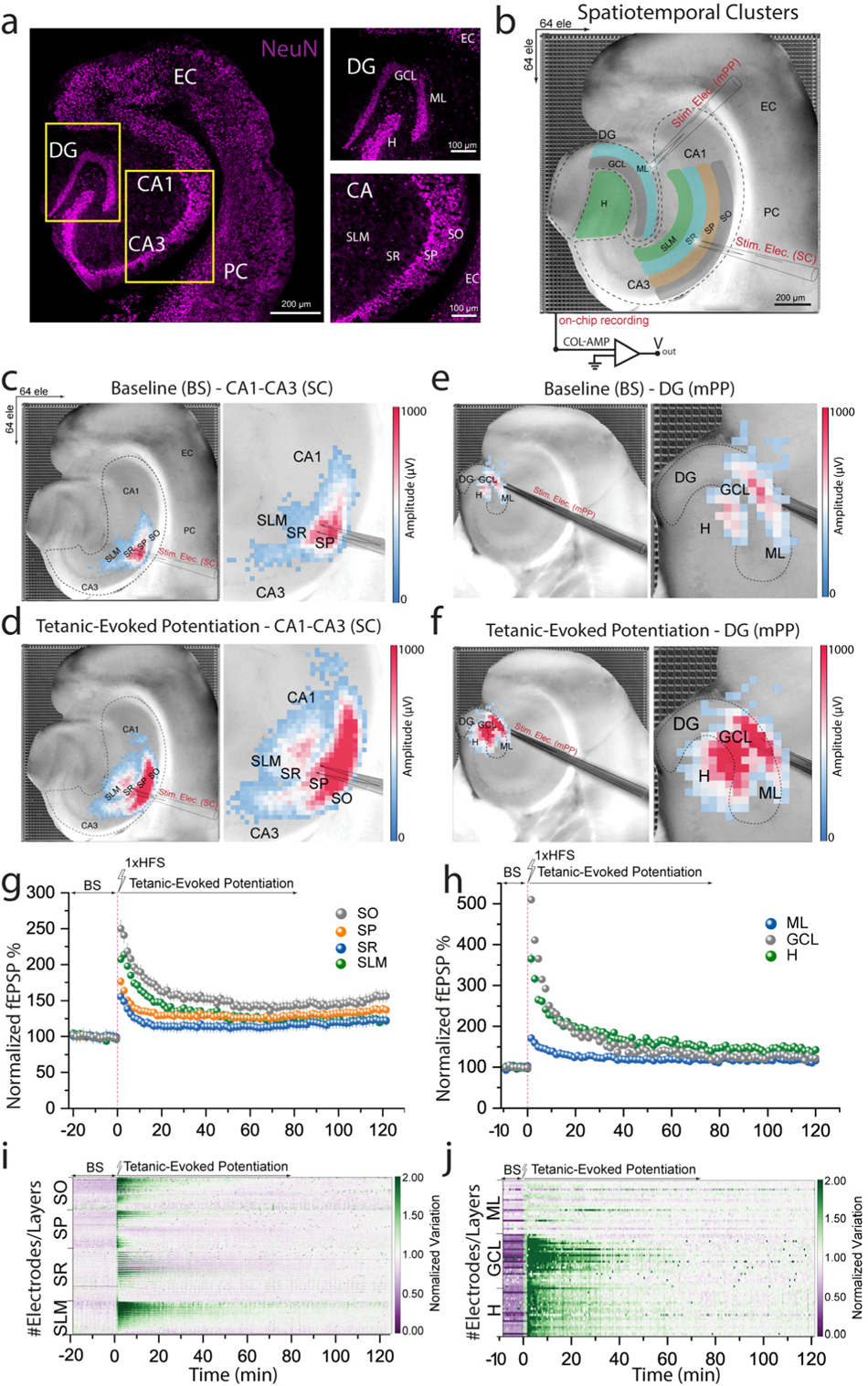
Detailed Mapping of Network-LTP-Induced Synaptic Changes in the Hippocampus. **a)** Fluorescence images of hippocampal subregions to identify distinct neuronal layers in the network**. b)** The platform overlaying functional bioelectrical readouts onto optical images of hippocampal subregions, mapping evoked synaptic activation in various neuronal populations across distinct hippocampal layers (SO, SP, SR, and SLM) in CA1-CA3 network and (ML, GCL, and hilus) in the DG. **c-f)** Pseudo-color maps showing dynamic representations of baseline and tetanic-evoked potentiation (LTP) for CA1-CA3 and DG networks, highlighting spatial heterogeneity and identifying the exact loci of synaptic strengthening. **g, h)** Quantification of normalized fEPSPs demonstrating biphasic LTP patterns characterized by immediate and persistent potentiation phases. **i, j)** Normalized voltage variation color-coded maps for each hippocampal sublayer, revealing layer-specific sensitivity to LTP induction and distinct voltage fluctuations during synaptic potentiation phases.

To examine the induction of network LTP and its effects on synaptic plasticity, we employed a high-frequency tetanic stimulation (HFS) protocol with a sequence of electrical pulses at 100 Hz in 1-sec duration. This aims to mimic the natural patterns of neuronal activity that lead to synaptic strengthening, ensuring robust LTP induction. Post-stimulation, we observed enhanced synaptic potentiation across the hippocampal network. The multisite recordings provided detailed insights into the temporal and spatial dependencies of LTP, revealing complex interactions between the stimulation patterns and LTP induction across different hippocampal regions.

Using pseudo-color maps, we dynamically represented baseline (BS) and tetanic-evoked potentiation (LTP) states for both CA1-CA3 and DG networks (**Figures 2c-f**). This visual quantification clearly distinguished between baseline and potentiated states, pinpointing the exact loci of synaptic strengthening within the network. It also highlighted the spatial heterogeneity of LTP effects, revealing previously inactive zones that showed significant strengthening post-LTP induction (**Figures 2d, f**).

We further quantified LTP phases based on normalized fEPSPs, defining initiation with 40% of baseline synaptic potential in the CA1-CA3 (**Figure 2g**) and DG networks (**Figure 2h**). Stimulation of SC and mPP fibers revealed a biphasic LTP pattern: immediate potentiation followed by persistent potentiation, which decayed during the late phase (up to 120 minutes). This biphasic pattern was consistent across all hippocampal subfields, demonstrating the robustness of our stimulation protocol. The quantification mapped spatially evoked responses and provided insights into the temporal dynamics of synaptic strengthening. We also computed the percentage of plasticity post-tetanic stimulation relative to baseline, revealing significant synaptic change dynamics at the network level: 82% ± 9.7 in SO, 47% ± 5.3 in SP, 31% ± 5.3 in SR, and 54% ± 3 in SLM within CA1-CA3; and 35% ± 5.6 in ML, 157% ± 8.2 in GCL, and 122% ± 8.4 in H within DG (*p < 10^-8^ Kolmogorov-Smirnov test*).

Next, we computed normalized voltage variation color-coded maps for each hippocampal sublayer (**Figures 2i, j)** to identify LTP-evoked response dependencies on spatial distribution. These maps revealed subtle voltage fluctuations during synaptic potentiation at the network level, showing distinct variations across different hippocampal layers. Notably, significant voltage changes were observed in SO and SLM of CA1-CA3 regions and GCL in DG throughout the potentiation phases, suggesting a layer-specific sensitivity to LTP induction, consistent with observations from small-scale electrophysiological recordings^31^. This analysis underscores the spatiotemporal complexity of synaptic modulation within the hippocampus and its implications for neuronal circuitry, memory formation, consolidation, and retrieval. It reflects the sophisticated mechanisms by which the brain processes and stores information through feature-coding cell assembly repertoire^14,32^.

Our findings support the concept that memory is dynamically shaped by neural activity patterns rather than being a static representation^33,34^. The EVOX platform, by combining multiple methodologies^35,36^ into a single, powerful tool, provides unparalleled resolution for mapping the entire hippocampal network. This opens new avenues for exploring how memories are encoded and manipulated within this crucial brain structure. By characterizing these sequential encoding patterns of network-wide LTP, we gain deeper insights into the neural basis of learning and memory.

Having characterized the sequential encoding patterns across the hippocampal network, we now delve into the specific synaptic signatures that emerge from these patterns, utilizing unsupervised classification to unravel the intricate waveform repertoires that define synaptic plasticity.

#### 2.3 Unveiling Synaptic Signatures through Unsupervised Classification of Waveform Repertoire

A critical question is how a detailed analysis of layer-based waveform signatures can enhance our understanding of synaptic plasticity and network dynamics in the hippocampus. EPSPs and population spikes (PS) are hallmark signatures of evoked neural activity and synaptic function within the hippocampus, typically studied through various conventional recording methodologies^12,37,38^. While these phenomena often serve as proxies for neuronal potentiation and collective neuronal firing, our approach moves beyond this binary framework. Our approach, however, utilizes the comprehensive multidimensional data from large-scale recordings to identify complex waveform signatures and their spatiotemporal characteristics, offering a deeper understanding of synaptic interactions in memory processes and neural circuit functionality^39^. This method enabled us to perform unsupervised analysis of numerous waveform patterns across different layers of the CA1-CA3 and DG networks, both at baseline and following tetanic stimulation.

Using principal component analysis (PCA) and k-means clustering^22^ (see Methods), we identified four distinct waveform classes within the CA1-CA3 network, corresponding to (SO, SP, SR, and SLM) (**Figures 3a-c**). Similarly, three unique evoked firing patterns were identified in the DG network, associated with (ML, GCL, and H) (**Figures 3e-g**). This spatially-resolved classification reveals how different hippocampal layers uniquely contribute to synaptic plasticity and functional reorganization.

**Figure 3.**
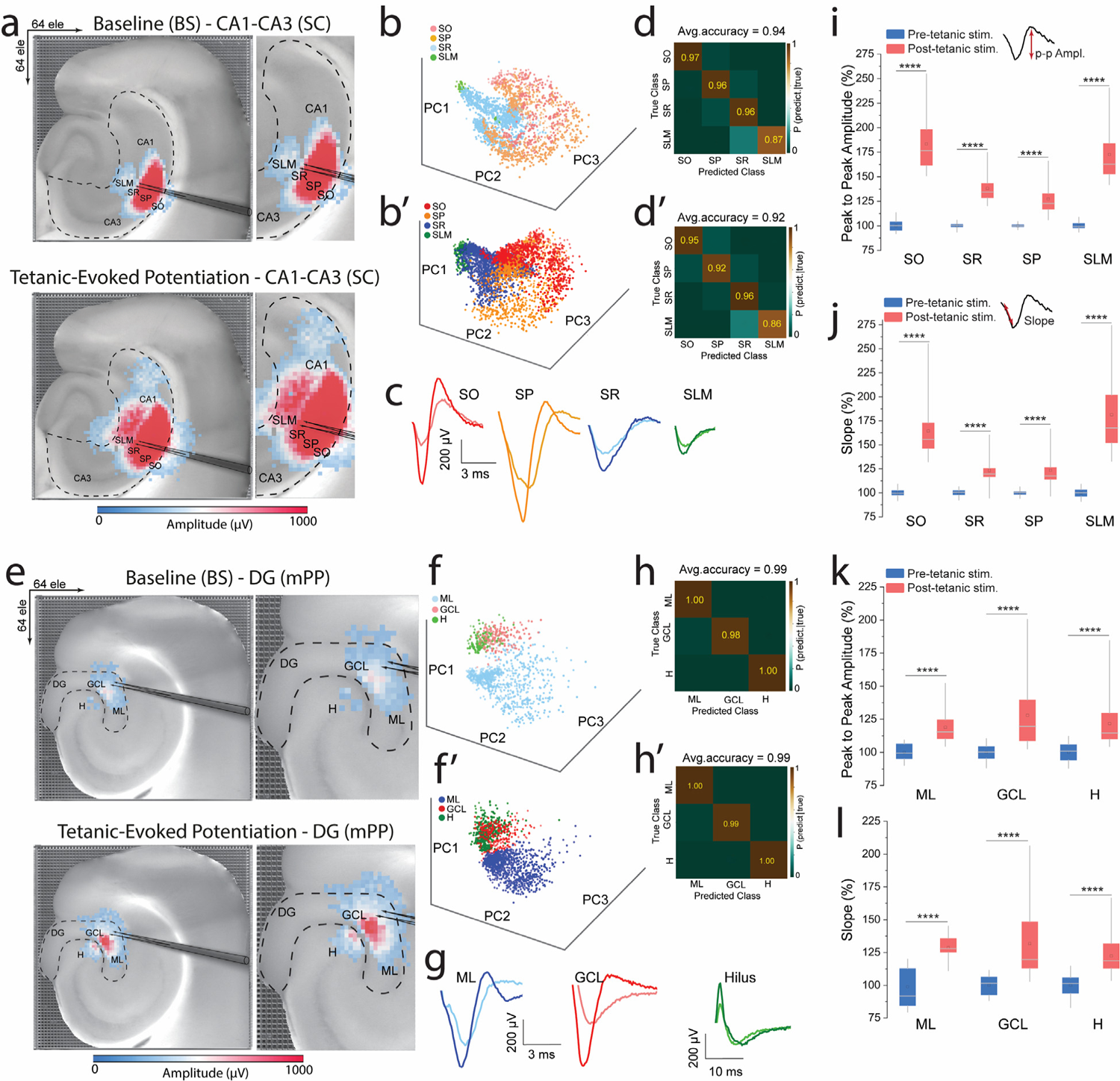
Classification and Analysis of Network-wide Evoked Synaptic Response Patterns. **a-c)** Principal component analysis (PCA) and k-means clustering identified four distinct waveform classes in the CA1-CA3 network layers (SO, SP, SR, and SLM) in baseline and tetanic-evoked potentiation phases. **e-g)** Similar classification in the DG network layers: molecular layer (ML, GCL, and hilus). **d, d’, h, h’)** Confusion matrices of K-means clustering accuracy scores, showing high precision in classifying and well-defined separation of evoked firing patterns before and after tetanic stimulation in CA1-CA3 and DG networks. **i-l)** Quantitative analysis of peak-to-peak amplitude and slope features of synaptic responses, demonstrating significant increases in these features across all hippocampal layers post-tetanic stimulation, indicating enhanced neural activation and synaptic plasticity.

Temporal analysis of these waveforms showed significant changes post-tetanic stimulation, indicating enhanced neural activation and improved discriminability of neural representations. The increased density and separation of data points in PCA maps post-stimulation (**Figures 3b, b’, f, f’**) highlights the selective strengthening of synaptic connections. Classification accuracy was high, with true probability scores of 94% ± 2.3 and 92.2% ± 2.2 for CA1-CA3 networks at baseline and post-tetanic phases, respectively. The DG network showed scores of 99.3% ± 0.6 and 99.6% ± 0.3 for the same phases, confirming the robustness of our classification method (**Figures 3d, d’, h, h’**).

We further analyzed the peak-to-peak amplitude and slope features of these waveforms during baseline and post-tetanic phases. This analysis revealed significant increases in both features across all hippocampal layers in response to tetanic stimulation (**Figures 3i-l**). These changes underscore the potential of potentiation-based waveform shapes as biomarkers for identifying layer-specific features within the hippocampal circuitry^40^. Additionally, the observed modifications in neural activity patterns induced by LTP emphasize the dynamic nature of synaptic plasticity and its critical role in information processing and memory formation.

Our findings underscore the importance of integrating spatial and temporal dimensions of firing signatures to fully understand synaptic plasticity and network reorganization associated with LTP^41^, which underlies activity-dependent modification of neural circuits and engrammed memory formation^42^. This granular, detailed analysis moves beyond traditional binary indicators, revealing the complex interplay of neuronal activity across various phases and layers of the hippocampus. The identified waveform features, informed by spatiotemporal dependencies, accentuate the capacity of EVOX to uncover novel biomarkers of synaptic performance and neural circuit integrity.

With a clearer understanding of the potentiation-based waveform signatures within hippocampal layers, our focus moves to mapping the spatiotemporal dynamics of evoked synaptic transmission, providing insight into how these dynamics facilitate complex learning and memory processes across the network

#### 2.4 Mapping Spatiotemporal Dynamics of Network-wide Evoked Transmission

Understanding the spatiotemporal dynamics of synaptic activation within the hippocampus is crucial for elucidating the mechanisms underlying learning and memory. Despite the established roles of neuronal firing patterns and synaptic plasticity in these processes, comprehensive mapping of these dynamics remains challenging^43,44^. Spike-timing-dependent plasticity (STDP) highlights the importance of temporal precision in learning mechanisms^45^. Additionally, the spatial organization of synaptic activation is vital for efficient information processing in the hippocampus, supporting spatial navigation and memory integration^46,47^. Despite advances, a complete understanding of these spatiotemporal dynamics poses significant challenges, representing a critical frontier in learning and memory research in the hippocampus.

To investigate these dynamics, we characterized extracellular synaptic responses by their post-stimulation latencies. We clustered peak time indices from processed evoked waveforms into initial, central, and terminal groups. We established correlations between the spatial distribution of synaptic events relative to the stimulating electrode and their temporal characteristics across CA1-CA3 and DG networks in both baseline and post-tetanic phases, as illustrated in the pseudo-color latency maps (**Figures 4a-d**). The time-based waveform clusters showed distinct timing of synaptic responses relative to the stimulation point (**Figures 4e-h**). We computed time delay dynamics and the distribution of activated electrodes in initial, central, and terminal clusters from baseline to post-tetanic phases. Notably, in the post-tetanic phase, we observed the emergence of new firing electrodes, indicating the activation of previously silent cell assemblies, reflecting significant restructuring of synaptic networks. This phase also showed a substantial reduction in synaptic response latency, suggesting increased synaptic efficacy across the hippocampal network.

**Figure 4.**
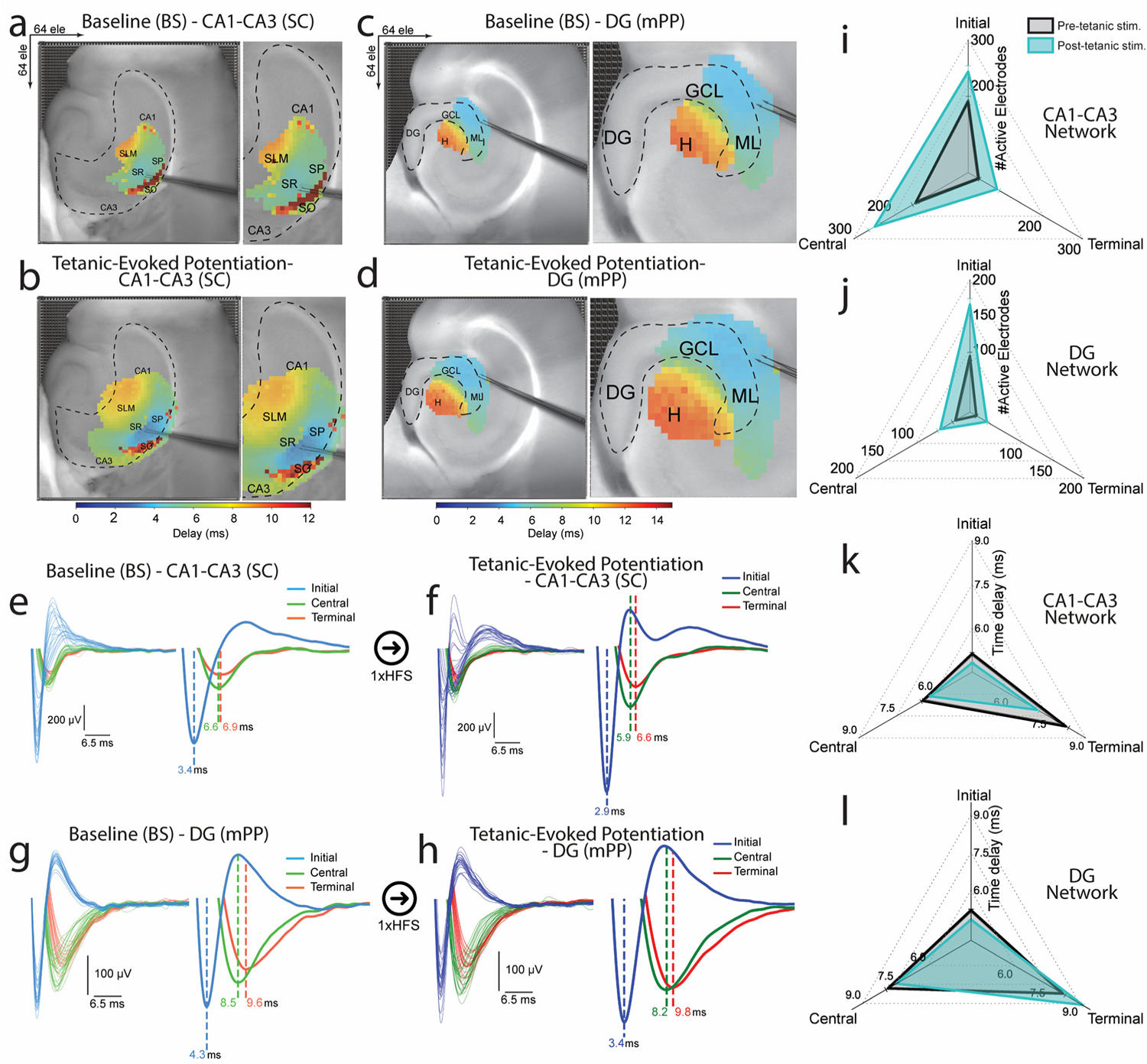
High-order Temporal and Spatial Dynamics of Network Synaptic Activation. **a-d)** Pseudo-color latency maps showing the spatial distribution of synaptic responses in the CA1-CA3 and DG networks at baseline and post-tetanic phases. These maps categorize synaptic events into initial, central, and terminal clusters based on their timing relative to the stimulating electrode. **e-h)** Temporal clustering of synaptic responses illustrates the precise timing of activation within hippocampal layers, highlighting significant changes in response patterns post-tetanic stimulation. **i-l)** Quantitative analysis reveals a substantial reduction in response latency and an increase in the number of active electrodes after tetanic stimulation, indicating enhanced synaptic efficiency and network restructuring. Notably, new firing electrodes emerge in previously silent regions, underscoring the dynamic reorganization of synaptic networks.

A quantitative analysis of the time delay and the number of active electrodes from multiple recordings confirmed a notable decrease in time delay (i.e., faster induction) and an expansion of the activated zone post-tetanically, indicating enhanced spatiotemporal efficacy of synaptic activity (**Figures 4i-l**). The activation of new firing electrodes, particularly in regions previously classified as silent or less active, marked a significant restructuring of synaptic networks. This was evident across CA1-CA3 (SO, SR, SP, and SLM) networks, with 1.5, 1.8, and 2.25-fold increases in initial, central, and terminal layers, respectively. Similarly, the DG (ML, GCL, and hilus) showed 1.8, 1.7, and 1.9-fold increases in initial, central, and terminal clusters, respectively, highlighting the potentiation of synaptic activity (**Figures 4i, j**). The normalized percent magnitude of the reduction in time delay from baseline to post-tetanic phases (**Figures 4k, l**) was 6% ± 0.3, 4.8% ± 0.4, and 13.8% ± 0.25 in initial, central, and terminal clusters in the CA1-CA3 network, and 7.3% ± 0.45, 4.8% ± 0.3 in initial and central clusters, but an increase in time delay of 10.8% ± 0.42 in terminal clusters of the DG network.

In the CA1-CA3 hippocampal network, our temporal mapping revealed the sequence of responses to tetanic stimulation, beginning with the SR, where direct excitatory inputs from the CA3 area are first integrated. This initial response is followed by activity in the SP, where these inputs are processed into neuronal outputs. Subsequently, the SO and SLM respond, reflecting their roles in modulating and integrating the network’s output through their complex inputs and indirect pathways. This unique orderly progression from initial integration (SR) through neuronal processing (SP) to modulation (SO and SLM) emphasizes the importance of understanding the sequence of synaptic activation and plasticity mechanisms in the hippocampus. Similarly, we mapped the temporal dynamics before and after tetanic stimulation in the DG layers, showing the sequence of synaptic activation and plasticity induction from the input layer (ML) through the processing layer (GCL) to the modulatory circuit (hilus). This progression underlines the complexity of information processing and synaptic modification in response to high-frequency stimulation within the hippocampus.

Our findings underscore the intricate interplay between the spatial positioning of synaptic activation and their temporal responsiveness to neural stimulation. This spatiotemporal framework enhances our understanding of synaptic organization in hippocampal layers and the dynamic processes of LTP and network adaptation. These insights highlight the hippocampus’s role in memory and learning by showing how changes in synaptic efficiency and new cell assembly activations contribute to cognitive functions. Our results may offer empirical evidence of how the hippocampus processes latent information, particularly in the DG–CA3 circuitry^48^, and how it dynamically reorganizes and encodes new information in response to external stimuli and experiences.

Building upon the detailed understanding of the spatiotemporal dynamics of synaptic transmission, the subsequent section will explore the contributions of mesoscale transmembrane generators to these dynamics, thereby advancing our comprehension of the complex synaptic interactions within the hippocampal network.

#### 2.5 Identifying Mesoscale Transmembrane Generators of Sequential Network-LTP Patterns

The relationship between electrical potential distributions and their corresponding sink and source currents in extracellular synaptic activation reveals essential insights into the principles of transmembrane generators and functional cell assemblies in the hippocampal circuit^49^. Variations in the magnitude and duration of these currents during synaptic events modulated by LTP provide a deeper understanding of synaptic transmission efficacy and the activation of complex neuronal assemblies with complex patterns coordinated across space and time. Recording methodologies with high spatiotemporal resolution are essential for clarifying these mesoscale synaptic dynamics and advancing our understanding of learning and memory processes.

In this study, we constructed two distinct spatiotemporal bidimensional maps—potential-based and kernel current source density (kCSD)^50^ —across a 64 x 64 grid that mirrors the HD-CMOS-MEA’s geometry. This approach allowed for the capture of dynamic shifts in synaptic activity and connectivity across various hippocampal subfields (SO, SP, SR, SLM) in the CA1-CA3 network, both under baseline conditions and after tetanic stimulation (**Figures 5a, b, f, g**). The potential-based maps facilitated detailed observations of extracellular potential signatures across different hippocampal layers, enhancing our understanding of synaptic dynamics and connectivity changes at the network level (**Figures 5c, h**).

**Figure 5.**
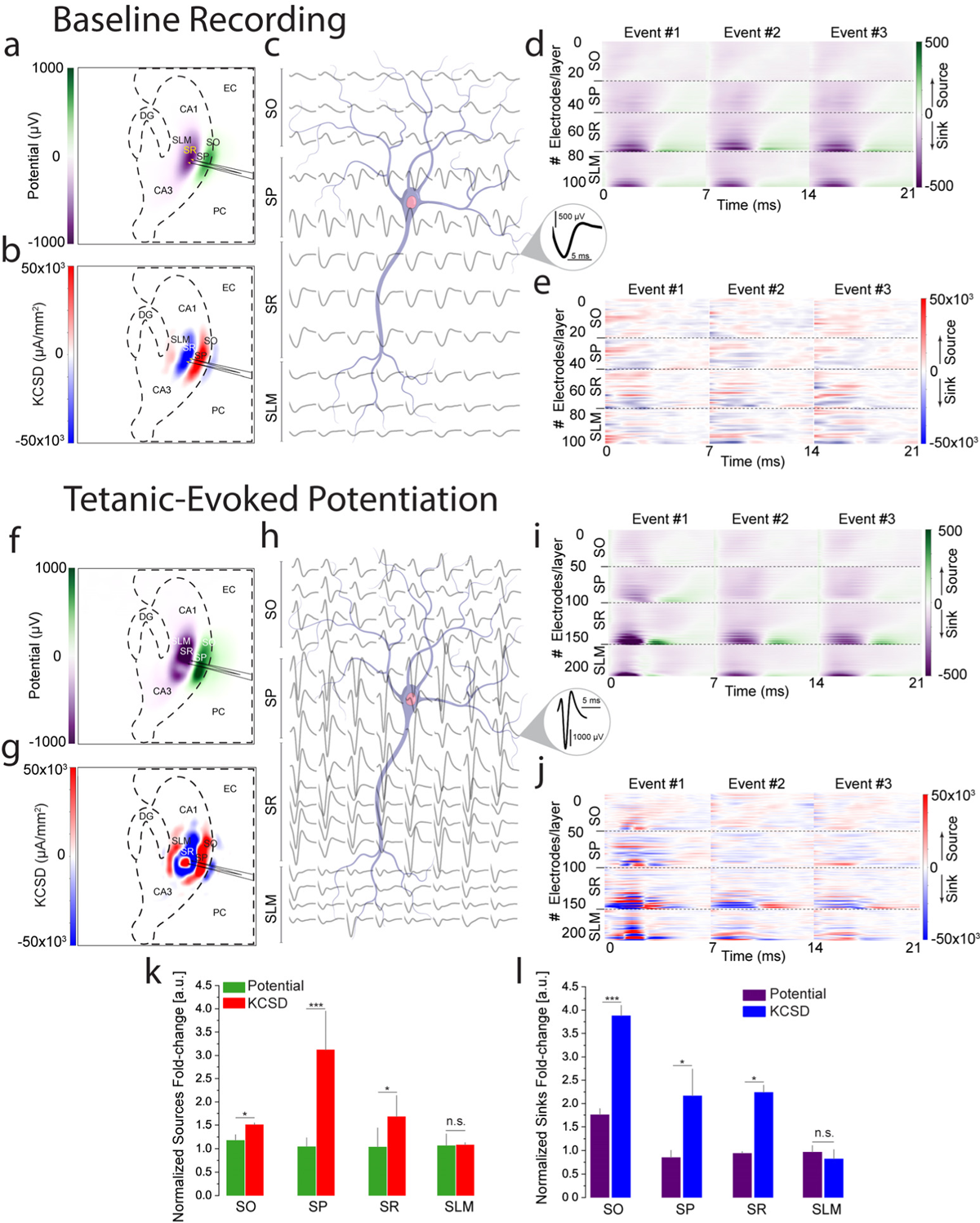
Mesoscale Mapping of Network Synaptic Activity and Connectivity Dynamics. **a, b, f, g)** Bidimensional maps showing potential-based and kernel current source density (kCSD) representations of synaptic activity in the CA1-CA3 network under baseline and post-tetanic stimulation conditions. **c, h)** Extracellular potential signatures across different hippocampal layers, illustrating network-level synaptic dynamics and connectivity changes from baseline to tetanic-evoked potentiation. **d, e, i, j)** Time-resolved synaptic activity maps, with the x-axis representing the timing of evoked events and the y-axis corresponding to firing electrodes within each layer. These maps provide a detailed view of sinks (associated with incoming EPSPs) and sources (related to synchronous neuronal firing). **k, l)** Comparison of quantified potential sources and sinks against those identified in the kCSD maps, highlighting the kCSD analysis’s superior precision in localizing synaptic events. This approach enhances our understanding of synaptic transmission and network dynamics in response to LTP, revealing intricate differences in sink and source values both at baseline and post-tetanic conditions.

Moreover, we organized the evoked synaptic data pre- and post-tetanic stimulation, aligning the x-axis with the timing of each evoked event and the y-axis with specific firing electrodes. This arrangement, coupled with computed potential-based and kCSD-derived source/sink generator data, provided a dynamic, spatial, and temporal view of synaptic activity (**Figures 5d, e, i, j**). This dual-mapping computational approach enabled a layered, time-resolved representation of synaptic patterns, highlighting sinks associated with incoming EPSPs and sources related to population spikes of synchronous neuronal firing.

Comparative analysis of quantified potential sources and sinks against those identified in the kCSD maps underscored pronounced differences. The potential-based analysis provided a general view of synaptic potentiation yet struggled to accurately resolve the spatial origin of synaptic events, which made it challenging to differentiate between EPSP and PS responses associated with precise sink/source generators. In contrast, the kCSD analysis offered a more precise localization of synaptic events, providing clearer insights into the underlying synaptic dynamics. Despite challenges in comparing these analyses due to their fundamentally different informational nature, their combined use is crucial for a comprehensive understanding of synaptic activity and connectivity. This nuanced view illustrates how LTP modifies the synaptic landscape, informing our understanding of the mechanisms underlying memory storage and retrieval within neural circuits.

These findings, consistently observed across multiple preparations, confirmed the detailed differences in sink and source values through potential computation and kCSD analysis in both baseline and post-tetanic conditions (**Figures 5k, l**). This comprehensive approach not only confirmed the consistency of our observations but also highlighted the complex nature of synaptic changes associated with network-wide LTP.

The detailed exploration of mesoscale transmembrane generators has underscored the capabilities of the EVOX platform in elucidating network-level synaptic potentiation and memory storage mechanisms. Leveraging these advanced insights, the subsequent section will focus on how aging modifies these synaptic processes, employing the EVOX platform to discern the variable impacts across different hippocampal layers.

#### 2.6 Network-resolved LTP and Synaptic Transmission in Aging Hippocampal Networks

Studying synaptic dynamics at the network level in aging circuits is essential for understanding how aging affects learning and memory. Fully grasping the influence of aging on hippocampal networks demands comprehensive, simultaneous monitoring of synaptic activity throughout the entire hippocampus, manifested through LTP induction. Methodological limitations have restricted a deeper insight into these age-related synaptic modifications.

Using the EVOX platform, we explored how aging alters synaptic function by examining layer-specific responses to LTP induction within the CA1-CA3 network. This approach allows for the detailed study of synaptic efficacy and neuronal sequence deactivation, contextualized by intrinsic aging of the circuitry. Upon establishing a stable baseline, a single HFS session triggered LTP across the entire network in aged mice. The fEPSP was quantified within the SP and SR layers of the CA1-CA3 network compared to the control group of young adult mice (**Figure 6a**). Notably, we observed a marked decline in synaptic plasticity in aged circuits compared to younger counterparts, with significant reductions in the fEPSP measurements: 38% ± 4.7 in SP and 21% ± 3.9 in SR, contrasted with 49.9% ± 5.9 and 24.8% ± 3.5, respectively, in young controls (*p < 0.01, Kolmogorov-Smirnov test*).

**Figure 6.**
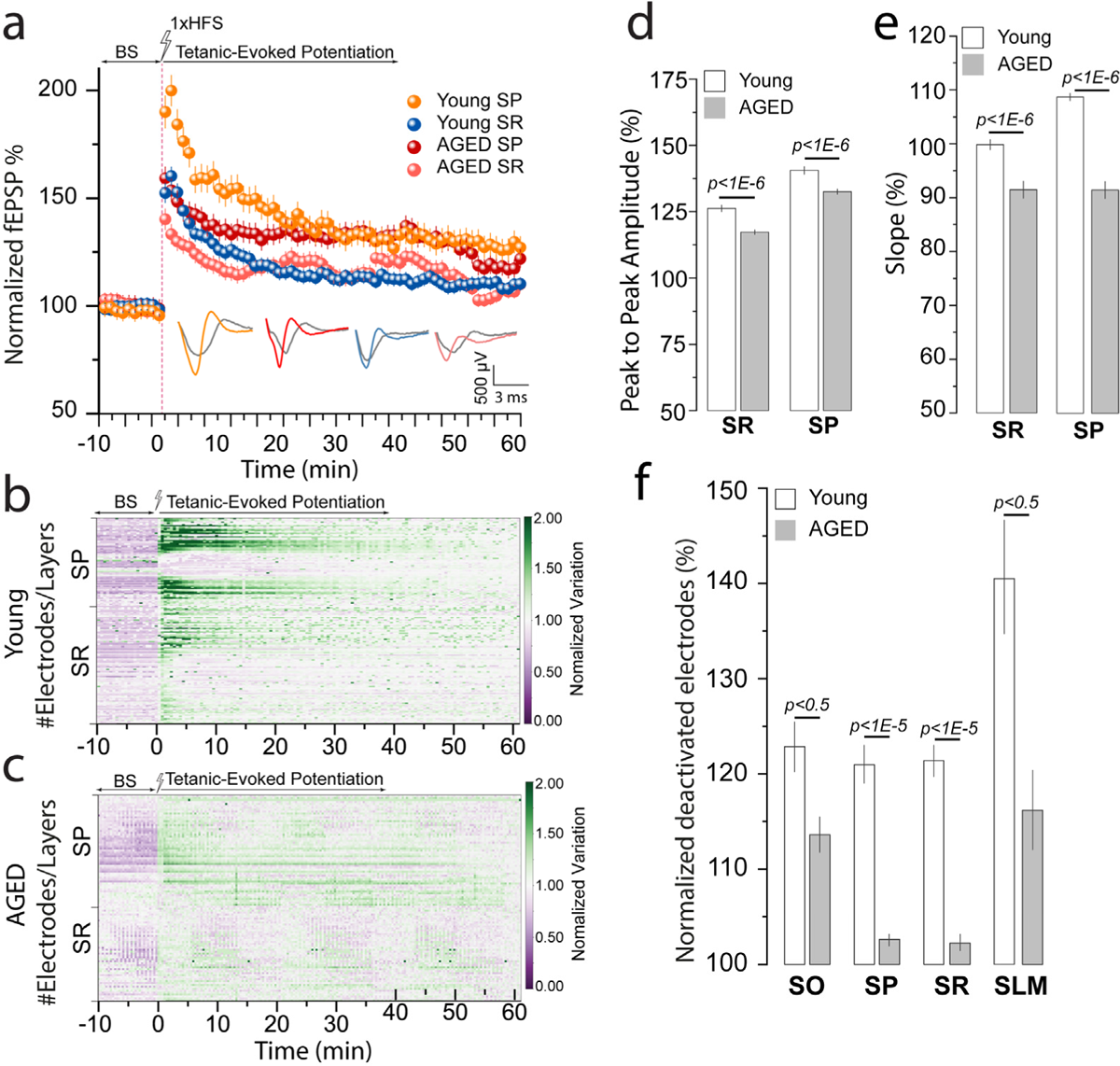
Impact of Aging on Network-Level Synaptic Plasticity in the Hippocampus. **a)** Comparison of normalized network-based fEPSPs in CA1-CA3 layers (SP and SR) between Aged and young standard mice, showing a significant decline in synaptic plasticity in the aged group. Also, this decline is identified in the layers-specific EPSP and PS patterns under the potentiation curve. **b, c)** Normalized voltage variation color-coded maps illustrating spatial distribution of LTP-evoked responses across hippocampal sublayers in Aged versus control young groups, highlighting intricate layer-specific quiescence and potentiation dependencies. **d, e**) Quantitative waveform analysis of synaptically evoked activity in SP and SR layers, revealing aging-related changes in amplitude and slope features during tetanic-evoked potentiation phases. **f)** Analysis of changes in deactivated firing associations from baseline to post-tetanic phases in the entire CA1-CA3 network, indicating a significant decrease in firing associations in the AGE group compared to SD young. These findings emphasize the influence of aging on network-level synaptic function and plasticity, offering insights into the dynamic reorganization of hippocampal circuits.

Further, our use of normalized voltage variation color-coded maps revealed how LTP-evoked responses depend on the spatial distribution across hippocampal sublayers SP and SR in the aged versus young adult groups (**Figures 6b, c**). These findings unveiled a distinct layer-potentiation dependency, showcasing how synaptic dynamic undergoes nuanced shifts across hippocampal sublayers—a testament to the layered complexity of synaptic integration and its susceptibility to aging-related modulations.

Moreover, a quantitative waveform analysis of synaptically evoked activity in layers SP and SR revealed critical insights into the amplitude and slope features of spatially identified responses in the tetanic-evoked potentiation phases (**Figures 6d, e**), highlighting the impact of aging on network synaptic functionality.

Our comprehensive examination also encompassed other layers in the CA1-CA3 network, where we identified changes in neuronal firing associations evidenced by a reduced count of active electrodes from baseline to post-tetanic phases (**Figure 6f**). These deactivated assemblies highlight a significant decline in synaptic connections within aged mice. Such changes underscore the effects of aging on the coordination among neuronal assemblies, crucial for maintaining strong synaptic transmission and efficient memory processing across the hippocampal layers. These findings provide vital insights into the age-related adaptations in network-based mechanisms of Hebbian learning and synaptic connectivit^51^.

In summary, our study using the EVOX platform sheds light on the landscape of synaptic plasticity within aging hippocampal networks and its implications for learning and memory. The observed dynamical changes in synaptic activation at the network level and the detailed characterization of age-related modifications provide a foundation for further studies into aging circuits. These insights enhance our understanding of the mechanisms underlying age-related cognitive decline and offer potential strategies for mitigating these effects, enriching our approach to neurodegenerative disease models and cognitive resilience enhancement in aged populations.

### 3 Discussion

Our platform marks a significant advancement in understanding the spatiotemporal dynamics of synaptic plasticity and LTP within the hippocampal network. Employing high-density microelectrode arrays, EVOX overcomes the limitations of traditional methods such as two-electrode recordings, conventional MEAs, and optical imaging methodologies. This advancement enables precise, label-free, comprehensive mapping of synaptic dynamics—essential for decoding the complex cellular interactions that underpin learning and memory processes. A fundamental strength of this study is its capacity to mimic natural neuronal activities at the network level through direct electrical tetanic stimulation, providing a controlled *ex vivo* environment that closely mirrors *in vivo* conditions. This sophisticated modeling of real-time synaptic dynamics across hippocampal layers offers unprecedented insights into the neural underpinnings of cognitive functions, substantially enriching our understanding of how synaptic activities facilitate memory processes^34,52^.

Our findings highlight how spatial and temporal dependencies critically influence synaptic efficacy, revealing intricate modulatory mechanisms that shape learning and memory. These insights affirm the significance of large-scale neural dynamics in orchestrating memory formation and retrieval processes, showcasing the dynamic nature of neural networks^53,54^.

We have elucidated the functional dynamics of the hippocampus, emphasizing the crucial role of sequential activation from the mPP to DG and from the SC to the CA regions. This supports the concept of the hippocampus as a dynamic indexer, intricately linking detailed memories across the cortex through well-coordinated cell assemblies^55^. Our observations on how various phases of network-LTP are vital for memory encoding in specific layers highlight the potential of EVOX to propel our comprehension of complex neural mechanisms.

The study also provides empirical evidence for representational drift, a mechanism suggesting the adaptability of memory systems essential for the ongoing updating and retrieval of memories^56^. This phenomenon could underlie the evolution of firing patterns of neuronal populations over time as a transition from baseline activity patterns to those induced post-tetanically. This adaptability is crucial for incorporating newly acquired information into long-term memory, enhancing the flexibility and resilience of memory networks^57^. Our findings also bolster the memory allocation hypothesis, demonstrating that highly excitable cell assemblies are preferentially recruited during memory encoding^58^. By mapping excitability shifts across hippocampal layers during different phases of LTP induction, we illustrate the dynamic selection of neuronal populations pivotal for memory formation. This supports the theory that memory encoding involves selective recruitment based on activity levels.

In terms of aging, our data reveal how synaptic plasticity is affected, with certain hippocampal layers showing decreased synaptic transmission and LTP induction in aged compared to younger counterparts. These age-related changes offer potential biomarkers for cognitive decline, providing insights into the synaptic foundations of aging within the hippocampus and implications for early detection of neurodegenerative conditions such as Alzheimer’s disease^29,59^.

The foundational research conducted with our platform has successfully highlighted key aspects of synaptic dynamics and plasticity, setting the groundwork for a range of exciting future investigations that extend beyond the initial scope of our study. These include exploring metaplasticity^60^, the interconnectedness of cell assemblies^61^, and the broader cellular mechanisms that underpin synaptic plasticity, such as NMDA receptor dynamics and the role of neuromodulators^12^. Future studies could bridge the gap between neuroscience and practical Brain-inspired technological applications, such as machine learning and neuromorphic computing, leveraging our findings to enhance the learning capabilities of artificial systems^62^. This comprehensive approach not only extends our initial findings but also paves the way for groundbreaking interdisciplinary research that merges neuroscience with technological innovations, setting a new course for future explorations that harness a deep understanding of neural dynamics, promising to revolutionize both theoretical and practical aspects of neuroscience.

### 4 Experimental Section

#### Animals and Housing Conditions

To investigate network synaptic transmission and LTP, experiments were conducted on C57BL/6J female mice aged 12 weeks (young adult group) and 56 weeks (aged group), obtained from Charles River Laboratories, Germany. All procedures complied with relevant European and national regulations (Tierschutzgesetz) and received approval from the local authority, Landesdirektion Sachsen (approval number 25–5131/476/14).

#### Acute Hippocampal Brain Slices Preparation

Mice were anesthetized with isoflurane before decapitation, and slices were prepared according to our previous studies^22–24^. The brain was carefully removed from the skull and placed in an ice-chilled high-sucrose cutting solution before slicing. The brain was securely placed in a custom-made agarose-based container and then affixed to the cutting plate. Horizontal slices of 300 μm thickness were prepared using Leica Vibratome VT1200S (Leica Microsystems, Germany). Slices were cut at 0–2 °C in a high sucrose artificial cerebro-spinal fluid (aCSF) solution saturated with 95% O_2_ and 5% CO_2_ (pH = 7.2–7.4) containing in mM: 250 Sucrose, 10 Glucose, 1.25 NaH_2_PO_4_, 24 NaHCO_3_, 2.5 KCl, 0.5 Ascorbic acid, 4 MgCl_2_, 1.2 MgSO_4_, 0.5 CaCl_2_. Next, hippocampal-cortical slices were incubated for 45 min at 32°C and then allowed to recover for at least 30 min at room temperature. Recording aCSF solution used during electrically evoked-synaptic response recordings contained in mM: 127 NaCl, 3.5 KCl, 1.25 NaH_2_PO_4_, 26 NaHCO_3_, 10 Glucose, 1 MgSO_4_, 2.5 CaCl_2_, and was saturated with 95% O_2_ and 5% CO_2_.

#### Extracellular Evoked-synaptic Responses and Network LTP in EVOX platform

The EVOX platform facilitated extracellular recordings using an HD-CMOS-MEA (3Brain AG, Switzerland) customized to accommodate our specific recording and stimulation protocol. This HD-CMOS-MEA consists of 4,096 electrodes, each separated by a 42 μm pitch covering an active sensing area of approximately 7 mm². To interface brain tissue slices with the electrodes, we employed a custom-designed platinum harp positioned directly above the tissue to ensure optimal contact. For sustained tissue viability and consistent experimental conditions, we implemented a temperature-regulated perfusion system. This system continuously supplied aCSF to the tissue-electrode interface at a flow rate of 4.5 mL/min, maintaining a constant temperature of 37°C throughout the experiments. All extracellular synaptic recordings were performed at 14 kHz/electrode sampling frequency. To facilitate detailed studies on large-scale evoked synaptic responses and LTP, we enhanced the EVOX platform with a precision zero-drift triple-axis micromanipulator system (SENSAPEX, Finland). Additionally, a bipolar electrode (70% Platinum, 30% iridium) was utilized (World Precision Instruments, Germany). The electrode featured a tip diameter of 3 μm, an outside diameter of 0.356 mm, and a length of 51 mm, with a tip separation of 125 μm at a nominal impedance of 2 MΩ, enabling precise selectivity of single and multiunit stimulation.

#### EVOX Stimulation-Recording Protocol

To generate sequential fEPSPs, the bipolar electrode was positioned either in the mPP of the DG or in the SR of the CA region to stimulate the SC pathway. After an accurate electrode placement into the tissue with our zero-drift manipulator, we applied a monophasic constant voltage pulse, with a pulse half-width ranging from 70 to 140 µs, and monitored the evoked responses. To calibrate the optimal stimulation intensity, we established an input/output curve by incrementally increasing the stimulation in steps of 10 µA from 20 to 130 µA at 30-second intervals, maintaining the same pulse half-width. The intensity that evoked 60% of the maximum fEPSP slope was identified and used for both baseline and tetanic-evoked potentiation phases. This selected intensity was automatically applied every 30 seconds until a stable baseline recording was achieved for 10-20 minutes. LTP was induced via a single high-frequency stimulation (HFS) train at 100 Hz, with a 10 ms interval and the same pulse half-width. We recorded evoked responses in the hippocampal regions up to 2 hours. Significant LTP was documented if the responses post-HFS during the tetanic-evoked potentiation phase were at least 40% greater than those during the BS phase^15^. After recording the evoked-synaptic responses, images of brain slices coupled to the HD-CMOS-MEA during network stimulation were captured by an optical modular stereomicroscope (Leica Microsystems, Germany). This also facilitated the analysis of the spatial organization and clusters of the tissue relative to the electrode layout.

#### Data Analysis

All analysis and algorithms used in this work were developed and implemented with custom-written Python scripts. Any package add-ons are cited accordingly.

#### Functional-Structural Clustering for Spatiotemporal Analysis

To analyze evoked-synaptic responses and LTP across specific hippocampal layers, we assigned firing electrodes to designated regions of the hippocampus. We generated topographical pseudo-color maps of large-scale evoked firing patterns from our recordings, which were superimposed on structural images of the hippocampus. Additionally, images captured with a light microscope were overlaid on the layout of the HD-CMOS-MEA to facilitate precise localization using Brainwave software (3Brain AG, Switzerland). Electrodes were systematically categorized into clusters corresponding to distinct structural landmarks within the hippocampal slice. These clusters encompassed six critical regions involved in hippocampal circuitry: the molecular layer (ML), granule cell layer (GCL), and hilus (H) within the DG, as well as the stratum oriens (SO), stratum pyramidale (SP), stratum radiatum (SR), and stratum lacunosum-moleculare (SLM) in the CA regions.

#### Automated Classification of Waveform Signatures

To systematically analyze the waveform signatures of evoked synaptic responses across distinct hippocampal layers, we employed an automated classification system integrating Principal Component Analysis (PCA) and k-means clustering^22,63^. The methodology was applied to elucidate the complex waveform shapes recorded from hippocampal circuits, specifically targeting responses from DG and CA layers following stimulation. Initially, PCA was employed to reduce the dimensionality of the data derived from sequential distinct stimulation events, capturing the primary features of the waveform shapes that signify different types of neural activity and focusing on the most informative aspects of the waveforms and their variance across samples. Subsequently, the k-means clustering algorithm was applied to these PCA-reduced data to categorize the waveforms into specific clusters^22^. Each cluster was associated with evoked responses from particular layers within the hippocampus: ML, GCL, and H in the DG and SO, SP, SR, and SLM in the CA regions. This classification facilitated the identification of distinct waveform patterns corresponding to different types of synaptic activity (i.e., EPSPs and PS) influenced by either mPP or SC pathway stimulations. To quantify the effectiveness of our classification, we computed the accuracy through the analysis of a confusion matrix. The average true positive rate was assessed by calculating the mean diagonal probability of the matrix, which compares the predicted class labels to the actual class labels. This probability ranges from 0 to 1, where a higher value indicates better accuracy and class specificity in the classification. The results from the confusion matrix were compelling, demonstrating well-defined separation and high accuracy in distinguishing between waveform classes associated with specific hippocampal layers. This procedure was partly implemented using the Scikit-learn 1.0.2: Machine Learning in Python and is available on GitHub (https://github.com/scikit-learn/scikit-learn/blob/main/sklearn/decomposition/_pca.py).

#### Framework for Temporal Clustering of Network-based Evoked Synaptic Responses

To analyze the temporal dynamics of evoked synaptic responses within the hippocampal layers, we employed a refined spatiotemporal clustering algorithm focusing on peak-time delays of electrically evoked synaptic responses. This approach enabled us to discern distinct clusters that reflect coordinated changes in synaptic timing, indicative of underlying synaptic plasticity and functional connectivity. It also allowed us to analyze evoked events from different experiments and animals and compare several slices’ responses regardless of the minor differences in the position of the external bipolar electrode to the mPP or SC region of stimulation interest. This involved applying a 4^th^-order Butterworth bandpass filter (1 – 500 Hz) to eliminate noise and artifact interference, ensuring the preservation of the waveform integrity. Following this, peak detection algorithms were employed to identify the exact timing of each peak within the waveform, which was crucial for the subsequent temporal clustering. We then categorized these identified peaks into three temporal clusters—initial, central, and terminal—based on their peak-time delays from the stimulus onset. This categorization was achieved through hierarchical clustering, which organized the peaks according to their temporal proximity to the stimulation. This allowed us to discern patterns in the timing of synaptic activations across different hippocampal layers. For validation, the robustness of the clustering process was assessed by analyzing the consistency of temporal groupings across multiple experimental sessions. Additionally, we examined the intra-cluster and inter-cluster variability to ensure the reliability of our temporal categorization. The final step involved visualizing and quantitatively analyzing these temporal clusters. We created pseudo-color latency maps to depict the distribution of synaptic events spatially and temporally, highlighting how different hippocampus regions responded over time to the stimulus. This visual representation and statistical analysis of changes in firing timings and electrode activations from baseline to post-tetanic phases provided deep insights into synaptic activation dynamics and synaptic plasticity’s underlying mechanisms. Through this methodology, we captured and elucidated the intricate temporal patterns of synaptic responses fundamental to understanding hippocampal function in learning and memory.

#### Kernel Current Source Density (kCSD) Analysis

To elucidate the sources and sinks of synaptic activity within the hippocampal layers using EVOX, we employed the kCSD analysis^25^, facilitated by the open-source kCSD-python package (https://github.com/Neuroinflab/kCSD-python/blob/master/kcsd/KCSD.py)^64^. This method is particularly well-suited for HD-CMOS-MEA because it can handle arbitrary electrode distributions, so it remains stable in case of contact malfunction. It uses regularization to reduce noise effects on analysis and accounts for differences in conductivity between tissue and the saline covering the brain slice^65^. The kCSD method employs a smoothing kernel to estimate the potential everywhere on the slice and a corresponding cross-kernel to move from the potential to the current source density^25^. These kernels encapsulate the conductive properties of the medium and geometry of the system.

The final output from this analysis is a two-dimensional spatial map of current source densities aligned with the electrode layout in distinct hippocampal layers, providing a detailed visualization of synaptic activity. This enhanced mapping capability of kCSD offers greater resolution and sensitivity in detecting and localizing subtle synaptic modifications than traditional potential-based analysis^64^. The clarity and precision in mapping synaptic events post-tetanic stimulation highlight dynamic synaptic interactions and connectivity changes. Utilizing kCSD in conjunction with the EVOX platform has markedly advanced our understanding of the intricate dynamics of synaptic responses, which is crucial for exploring the mechanisms underlying neural circuit functionality and synaptic plasticity.

#### Immunofluorescence Protocol

Fixed mouse brains were sectioned horizontally at 14 µm and processed for immunofluorescence using established protocols^66^. The cryosections were first rehydrated in 1× PBS and then permeabilized progressively with Triton X-100, decreasing concentrations from 0.3% to 0.1% in PBS (PBST). Antigen retrieval was performed by heating the sections in 10 mM citric acid (pH 6.0) at 95°C for 10 minutes, followed by cooling for 20-30 minutes and extensive washing in 0.1% PBST at room temperature. The sections were blocked in 0.1% PBST containing 5% normal goat serum for 1 hour at room temperature. Subsequently, they were incubated with primary antibodies diluted 1:1000 in the blocking solution overnight (14-16 hours) at 4°C in darkness. After several washes, the sections were incubated with secondary antibodies and diluted 1:1000 in blocking solution for 2 hours at room temperature. Following additional washing steps in 0.1% PBST and then 1× PBS, the sections were stained with Hoechst (1:1000 dilution from a 10 mg/ml stock solution, Thermo Fisher) for 30 minutes in darkness, washed thoroughly in 1× PBS, mounted with Fluoromount-G (Invitrogen, Germany), and left to air-dry overnight in darkness. The preparations were then sealed with nail polish (Electron Microscopy Sciences, Germany) and examined under an LSM 980 Airyscan 2 microscope (Zeiss, Germany). For immunolabeling, the following primary antibodies were used: Tuj1 (rabbit, Synaptic Systems, 302302, 1:500), Map2 (guinea pig, Synaptic Systems, 188006, 1:250), and NeuN (chicken, Synaptic Systems, 266006, 1:200). All primary antibodies underwent antigen retrieval to ensure optimal staining.

#### Statistical Analysis

All statistical analyses were performed with Python and Originlab 2024. All data in this work were expressed as the mean ± standard error of the mean (SEM). All box charts are determined by the 25th-75th percentiles and the whiskers by the 5th-95th percentiles and lengths within the Interquartile range (1.5 IQR). Also, the lines show the median and the squares for the mean values. Differences between groups were examined for statistical significance, where appropriate, using the Kolmogorov-Smirnov test, one-way analysis of variance (ANOVA), followed by Tukey’s posthoc testing. P-value < 0.05 is considered significant, and n.s. indicated non-significant.

## Authors Contributions

S.K. performed experiments, analyzed data, and generated the figures. X.H. wrote the code and analyzed the data. B.A.E. performed experiments. W.S. and D.W. helped with the kCSD analysis and discussed the study results. G.K. supported the interpretation of results and contributed to co-funding. H.A. conceptualized, planned, and supervised the project, designed and performed experiments, developed computational tools, and finalized the figures. H.A. coordinated the study and wrote the manuscript. All authors reviewed and approved the final manuscript.

## Acknowledgments

This study was financed from basic institutional funds (DZNE). We would like to thank Dr. Detlef Balschun (KU Leuven), Dr. Alessandro Maccione (3Brain AG), and Dr. Alexander Garthe (DZNE) for their insightful comments on the manuscript and the fruitful discussion. We would also like to acknowledge the support of the platform for behavioral animal testing at the DZNE-Dresden (Anne Karasinsky, Sandra Günther, and Jens Bergmann).

